# TWAS-CTL: A robust and efficient method for multi-tissue transcriptome-wide association studies using cross-tissue learners

**DOI:** 10.1101/2025.10.25.684514

**Authors:** Md Mutasim Billah, Hairong Wei, Fengzhu Sun, Kui Zhang

**Affiliations:** Department of Mathematics, Statistics, and Computer Science, Macalester College, Saint Paul, MN, USA; Department of Mathematical Sciences, Michigan Technological University, Houghton, MI, USA; Department of Quantitative and Computational Biology, University of Southern California, Los Angeles, CA, USA

## Abstract

The advent of transcriptome-wide association studies (TWAS) has expanded the classical genome-wide association study (GWAS) framework by integrating gene expression with genetic variation to identify trait-associated variants. While multi-tissue TWAS approaches improve statistical power over single-tissue models, existing methods often lose information during result aggregation and require intensive computation. Here, we present TWAS-CTL (Cross-Tissue Learner), a novel framework that leverages heterogeneous gene expression across tissues by adaptively reweighting and optimizing multiple single-tissue learners. Simulations demonstrate that TWAS-CTL achieves higher statistical power than the leading method, UTMOST, while maintaining proper type I error control and reducing computational time by over half. When applied to the analysis of the Genetics of Kidneys in Diabetes (GoKinD) cohort, we observed that TWAS-CTL identified more susceptible genes associated with diabetes than both UTMOST and PrediXcan, another widely used method in TWAS. These results establish TWAS-CTL as a powerful and efficient tool for cross-tissue gene expression analysis, capable of integrating heterogeneous gene-trait associations to advance genetic discovery.

## INTRODUCTION

Despite more than a decade of genome wide association studies (GWAS), which have now identified a large number of genetic variants that are associated with complex human traits and diseases (Tam et al., 2019; Visscher et al., 2017; Vujkovic et al., 2020), the biological mechanisms that link DNA sequence variation to clinical phenotypes remain largely unresolved. Many of these susceptibility variants exert their effects not by altering protein-coding sequences directly, but by modulating gene expression and transcriptional activity - an insight supported by numerous studies of expression quantitative trait loci (eQTLs) (Albert & Kruglyak, 2015; Tuuli Lappalainen et al., 2013; Zhang et al., 2015). However, measuring associating single-nucleotide polymorphisms (SNPs) with the traits of interest in cohorts large enough to detect modest effects is impractical (Spencer et al., 2009) because of the specimen availability and cost of measuring gene expression (Gusev et al., 2016). Early gene-based frameworks, as described by (Gamazon et al., 2015), (Zhu et al., 2016) and (Gusev et al., 2016) recognized this hurdle and introduced transcriptome-wide association studies (TWAS). TWAS leverages the relationship between genetic variants and gene expression, as characterized in reference transcriptome panels such as GTEx (Consortium et al., 2020) to impute genetically regulated expression in large GWAS cohorts. In practice, cis-acting SNP weights derived from the reference panel are used to predict gene expression levels across individuals with available genotype data, after which these imputed expression values are tested for association with phenotypic traits. This framework effectively bridges the gap between genetic variation and complex traits through the mediating layer of gene expression. The implementation of TWAS has substantially advanced our understanding of the genetic architecture underlying numerous complex diseases and traits (Barbeira et al., 2018; Nicholas Mancuso et al., 2017).

The latest GTEx release, together with the Depression Genes and Networks (DGN) study and other large consortia, has generated comprehensive multi-tissue transcriptomic resources, profiling dozens of tissues across hundreds of individuals and enabling unprecedented cross-tissue eQTL mapping (Saha et al., 2017; Yang et al., 2017). However, most current TWAS implementations train independent models for each tissue. This single-tissue strategy overlooks shared regulatory mechanisms across tissues, inflates multiple testing burdens, and may yield spurious associations in tissues that are not biologically relevant to the trait of interest (Liu et al., 2017; Pejman et al., 2017; Wainberg et al., 2017). By contrast, multi-tissue methodologies have demonstrated markedly improved statistical power in eQTL discovery (Flutre et al., 2013; Li et al., 2018; Sul et al., 2013), suggesting that a unified cross-tissue framework could similarly enhance the performance of TWAS. Building on this idea, the Unified Test for Molecular Signatures (UTMOST) was developed as a cross-tissue TWAS framework designed to improve gene expression prediction through a sparse group LASSO penalty (Hu et al., 2019). While UTMOST assumes that SNP effects are shared among tissues, it models these effects independently and therefore fails to fully capture inter-tissue similarity (Zhou et al., 2020). Yet numerous studies have shown that eQTL effects are frequently shared across functionally related tissues (Consortium et al., 2015; Pejman et al., 2017). Ignoring such cross-tissue similarity can lead to reduced statistical power, particularly when gene expression regulation is coordinated across multiple tissues. For example, tissues such as the pancreas, liver, brain, and blood exhibit extensive overlap in cis-regulatory architecture, while also maintaining distinct transcriptional programs that modulate disease susceptibility in tissue-specific ways. Existing multi-tissue TWAS approaches typically adopt one of two strategies: they either build separate gene expression predictors for each tissue or aggregate single-tissue association statistics post hoc. Both strategies fail to account for the graded continuum of regulatory sharing and divergence across tissues- that is, the spectrum spanning from regulatory homogeneity to strong tissue-specific heterogeneity. By overlooking this nuanced cross-tissue structure, current methods risk diminished statistical power and potential misattribution of association signals to irrelevant tissues.

To address the limitations of existing multi-tissue TWAS methodologies, we introduce Transcriptome-Wide Association Study- Cross-Tissue Learner (TWAS-CTL), a novel statistical framework that explicitly incorporates both tissue homogeneity and heterogeneity into model construction process. The framework operates in two stages. In the first stage, a single-tissue learner (STL) is trained independently for each tissue using any predictive algorithm capable of modeling the effects of cis-SNPs on gene expression (e.g., elastic net or other machine learning models). This process yields a set of tissue-specific expression predictors. In the second stage, each STL is applied to predict gene expression in all other tissues. A utility-based weighting function (Patil & Parmigiani, 2018) is then used to quantify how well each model generalizes across tissues. Tissues whose predictors transfer effectively to other tissues receive higher weights, while those exhibiting highly tissue-specific regulation are down-weighted. Tissues whose predictors generalize well across the tissue panel were assigned higher weights, while those with highly tissue-specific regulatory patterns were down-weighted accordingly. This utility-driven weighting scheme serves as an empirical measure of cross-tissue regulatory similarity, enabling TWAS-CTL to exploit shared genetic architecture while preserving sensitivity to tissue-specific effects. The design of TWAS-CTL provides two key advantages. First, it is learner-agnostic - any statistical, machine learning, or deep learning model capable of mapping cis-SNPs to expression can be incorporated as the STL without modification to the overall framework. Second, it is utility-modular - the weighting function can be flexibly defined based on evaluation metrics such as mean squared error, cross-validated correlation, or other customized criteria, allowing adaptation to the characteristics of the underlying genetic architecture and sample size. We comprehensively evaluated TWAS-CTL through extensive simulation studies encompassing a wide range of sample sizes, eQTL replication rates, and degrees of tissue heterogeneity. The results demonstrated that TWAS-CTL provides robust control of the nominal type I error rate while consistently achieving equal or superior statistical power compared to UTMOST. Finally, we applied TWAS-CTL to real-world data from the Genetics of Kidneys in Diabetes (GoKinD) cohort - a case-control study of type 1 diabetes. In this empirical setting, TWAS-CTL outperformed both PrediXcan and UTMOST, identifying a larger number of trait-associated genes and yielding stronger association signals, thereby highlighting its effectiveness and robustness as a novel framework for multi-tissue TWAS.

## MATERIAL AND METHODS

### Data Pre-processing and Quality Control

We used the GTEx (Genotype-Tissue Expression) V8 gene expression and genotype data as the reference panel for all analyses. Open-access gene expression data were downloaded from GTEx Analysis Release V8. Of the 838 donors, 715 (85.3%) were European American, 103 (12.3%) African American, and 12 (1.4%) Asian American; 16 individuals (1.9%) reported Hispanic or Latino ethnicity; The cohort was 66.4% male (n=557) and 33.5% female (n=281) (Consortium et al., 2020). Previous cross-tissue TWAS frameworks, including UTMOST, have employed RPKM (Reads Per Kilobase of transcript per Million mapped reads; GTEx V6) and TPM (Transcripts Per Million; GTEx V7) as normalization strategies for gene expression. However, both RPKM and TPM are optimized for within-sample normalization and are not appropriate for between-sample or cross-tissue comparisons, due to their inability to account for library size and compositional biases. Recent work (Johnson & Krishnan, 2022) indicates CTF (counts adjusted with timed mean of M levels factors) and CUF (count adjusted with upper quartile factors) are the best between-sample normalization methods. In our work, we downloaded gene read counts by tissue data (GTEx V8) for 49 tissues and then performed between-sample normalization. To be more specific, we used count adjusted with upper quartile factors (CUF). Additionally, we obtained access to the protected genotype data through authorized application via dbGaP. Following normalization, we further refined the gene expression data to remove known sources of confounding variation. Specifically, we regressed out the effects of biological sex, the top three genotype principal components, sequencing platform, and top PEER (Probabilistic Estimation of Expression Residuals) factors, in accordance with best practices for eQTL and TWAS analyses.

To get access to the protected genotype data we submitted an application through the NIH database of dbGaP for the GTEx protected access data with an accession id of phs000424.v8.p2, available at https://www.ncbi.nlm.nih.gov/projects/gap/cgi-bin/study.cgi?study_id=phs000424.v8.p2. It is crucial to perform quality control of the genotype data. We used PLINK version 1.9 beta, a publicly available, open-access toolkit for whole-genome association studies (Purcell et al., 2007). SNP-level missingness refers to the proportion of individuals lacking genotype data for a given SNP. High levels of missingness at certain SNPs can introduce bias into the analysis and are typically addressed during quality control procedures (Marees et al., 2018). Another key component of quality control is testing for adherence to Hardy-Weinberg equilibrium (HWE), which assesses whether the observed genotype frequencies at a given SNP deviate from those expected under assumptions of random mating in a large, stable population (Kalogeropoulou et al. 2009). In large, randomly mating populations that are not influenced by evolutionary or demographic forces altering allele frequencies, the genotype distribution for a particular marker is expected to adhere to the Hardy-Weinberg Equilibrium (Hosking et al., 2004). We removed the SNPs with a missing rate of more than 10%, SNPs that violate the HWE (p-value < 0.05), and the samples with a missing rate of more than 10% (Zhou et al., 2020). The individual heterozygosity rate, defined as the ratio of heterozygous genotypes per individual, functions as an essential quality control parameter in GWAS. Increased heterozygosity rates may signify sample contamination, whereas decreased rates may imply inbreeding. Individuals with heterozygosity rates that considerably differ from the mean, generally above three standard deviations, are routinely excluded (Marees et al., 2018). We removed 5 samples with the highest individual-heterozygosity rate and 5 samples with the lowest individual-heterozygosity rate- along with the PLINK, git bash was also used in this step (https://git-scm.com/downloads/win). Linkage disequilibrium (LD) refers to the non-independent association of alleles at different genetic loci. LD pruning entails the elimination of SNPs exhibiting substantial linkage disequilibrium to preserve a collection of relatively independent variants, thus diminishing redundancy and enhancing the efficacy of association studies. The choice of the r^2^ threshold for LD pruning is somewhat arbitrary and depends on the specific goals of the analysis. A common threshold is of 𝑟^!^ = 0.9 (Zhou et al., 2020) was used here. Minor allele frequency (MAF) represents how often the less prevalent allele appears within a given population. A SNP with a low MAF is less informative because it is less likely to be associated with a trait or disease. In addition, low frequency alleles are more likely to be affected by genetic drift, which can result in spurious associations. For these reasons, SNPs with low MAF are often excluded from genetic association studies. The cutoff of 0.05 was used, but it may vary depending on the specific study design and population being studied (Zhou et al., 2020). For the GTEx v8 genotype data (all chromosomes), after applying quality-control filters like SNPs with individual-level missingness, Hardy-Weinberg equilibrium, pruning out correlated, MAF threshold, 2645120 SNPs and 828 samples remain for downstream analysis.

### TWAS-CTL

The schematic depicts the logic of a modern transcriptome-wide association study (TWAS) as a two-panel workflow that links cis-regulatory variation to complex-trait risk in **Figure 1**. At the top of the schematic is the reference panel (e.g., GTEx), in which both genotype data (M SNPs across *n* individuals) and RNA-sequencing data are available across multiple tissues. In each tissue, an imputation model is trained to map local SNP variation to gene expression. The estimated weights from these models are then applied to the GWAS panel (*M* SNPs across *n*_1_ individuals). For each GWAS participant and for each tissue, the SNP weights yield a vector of imputed expression. Because the same genetic predictors are used across tissues, imputation propagates the cis-regulatory architecture learned in GTEx into the much larger disease cohort. TWAS then used the GWAS genotypes and the imputed expression profiles to regress the trait of interest (phenotype) on the predicted transcriptome to obtain the gene-trait association. Collectively, these steps constitute the foundation of contemporary multi-tissue TWAS: learning cis regulation where expression is available, projecting them onto large GWAS cohorts, and integrating the resulting signals to illuminate the genes - and the tissues - most plausibly involved in complex disease etiology.

**Figure 1.**
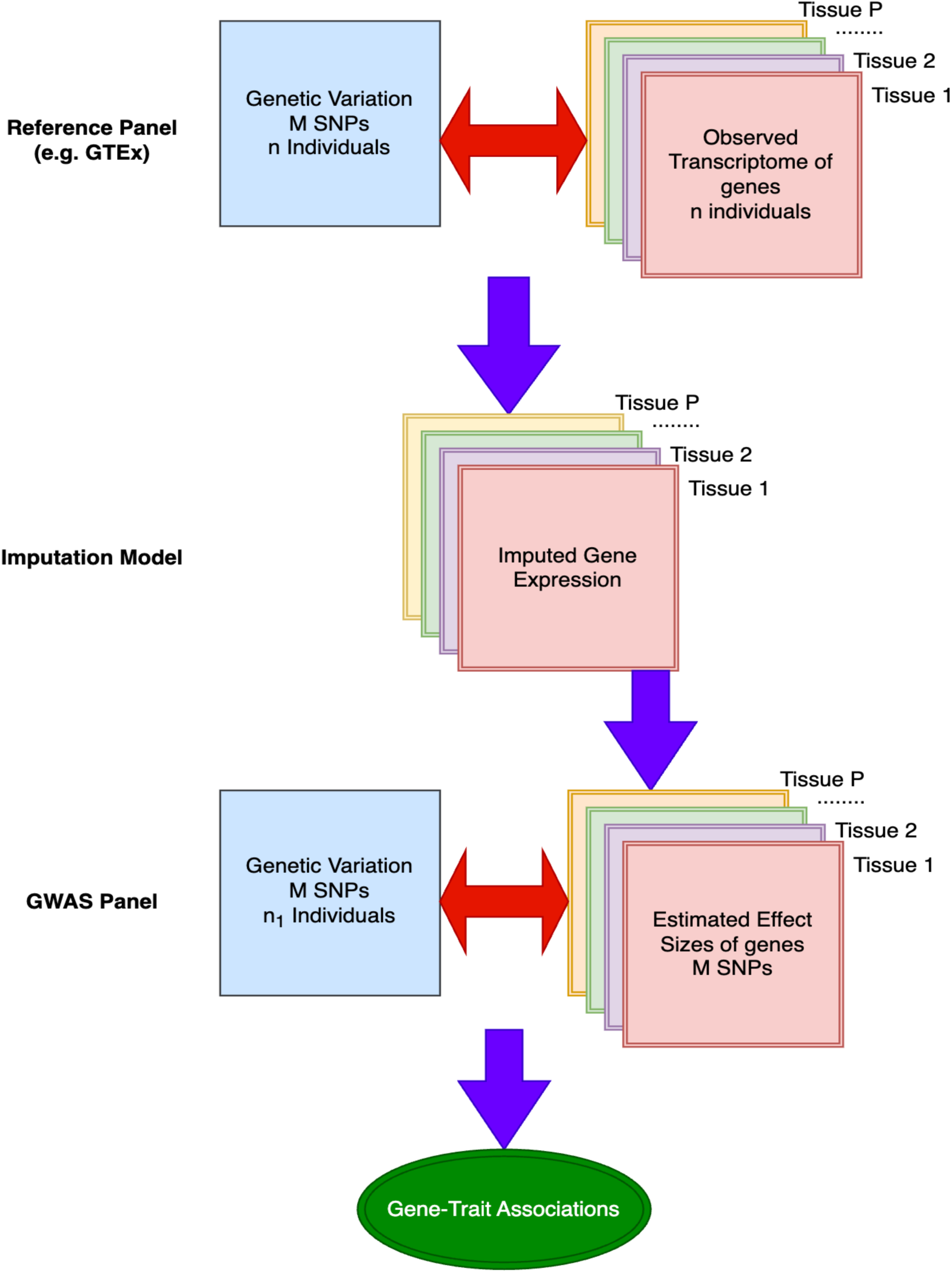
The Overview of Transcriptome-wide Association Studies (TWAS).

### Multi-tissue TWAS-CTL

The overall architecture of the TWAS-CTL (Cross-Tissue Learner) is illustrated in **Figure 2**. For each gene of interest, we begin by assembling expression measurements across P tissues alongside corresponding genotype data from a reference panel such as GTEx. In the first stage, we train one or more Single-Tissue Learners (STLs) independently for each tissue. Each STL - whether it be a penalized regression model, a tree-based learner, or another algorithm - learns the relationship between genotypes and gene expression within its tissue and produces a predicted expression profile. In the second stage, a bespoke weighting function evaluates the predictive performance of every tissue–model combination and assigns higher weights to those with stronger, more reliable signals while down-weighting noisy or uninformative contributions (Patil & Parmigiani, 2018). Finally, CTL aggregates the weighted STL predictions into a unified cross-tissue imputed expression, thereby leveraging both tissue-specific and shared genetic influences to enhance imputation accuracy and robustness compared to single-tissue or single-model approaches.

**Figure 2.**
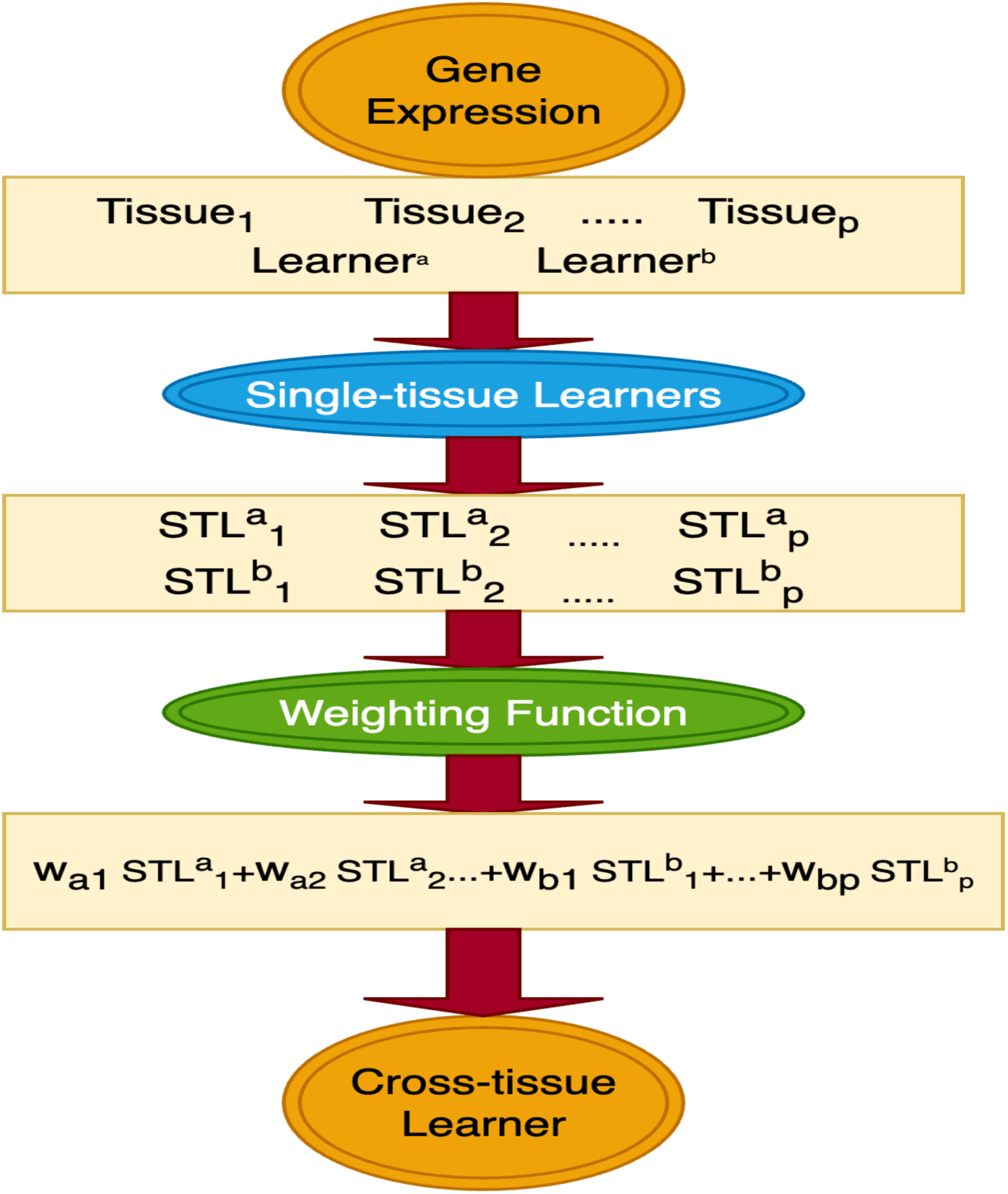
The mechanism of TWAS-CTL (cross-tissue learner in transcriptome-wide association studies).

For a given gene, gene expression values of n individuals in tissue j using learner l is (STL):

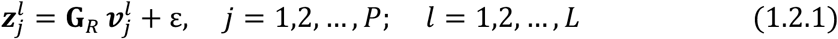

where 𝐆_𝑹_ is a 𝑛 by *M* matrix, representing the genotype of n individuals across *M* SNPs from the reference panel and 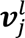 is a column vector with 𝑀 elements, representing the weights of *M* SNPs for j-th tissue.

The predicted expression in the j-th tissue, using learner l, is:

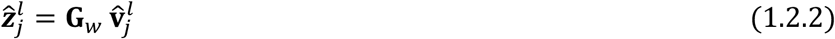

where, **G**_w_ is a *n*_1_ by *M* matrix, representing the genotype of *n*_1_ individuals across *M* SNPs from the GWAS.

Finally, for a given gene, the cross-tissue combined predicted gene expression (CTL) is:

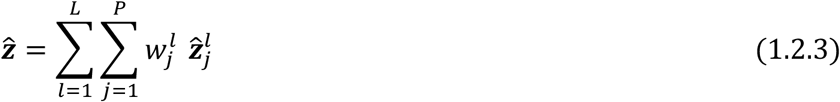

Here 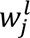 is the weighting function, which has been designed in such a way that it assigns a weight of zero to the worst STL, shifts losses for all other STL’s accordingly, and hence normalizes the weights. 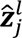 is being derived using equation (1.2.2).

We quantify cross-tissue predictive accuracy using a utility function: 𝑈 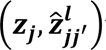 Awhere 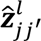 includes the predictions obtained using model 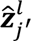, trained in tissue *j*’, on the prediction profiles in tissue *j*, using learner l.

For each learner, l, we construct a 𝑝 by 𝑝 matrix, 𝑿*^l^*, with entries:

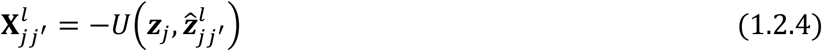

where the rows of 𝑿*^l^* represent how each 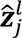 performs across all tissues.

For each learner l, we derive its weights by first summarizing the row j of 𝑿*^l^* by the mean of off-diagonal entries (omitting the resubstituting error 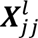)—

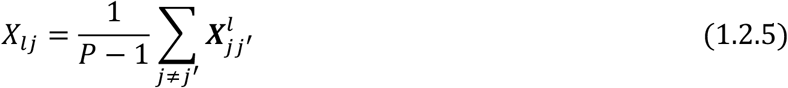

Alternative summary measures may also be applied; for instance, one could take the square root of each off-diagonal value before averaging.

Finally, the CTL weights can be found as:

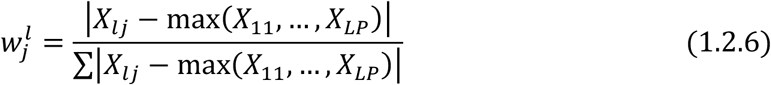

Under this scheme, the lowest-performing STL is assigned a weight of zero, the loss values for the remaining STLs are adjusted relative to that baseline, and the resulting weights are then normalized.

In this work, we used the following formula:

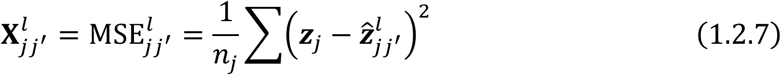

as the utility function and the square-root of the average off-diagonal mean-square error -

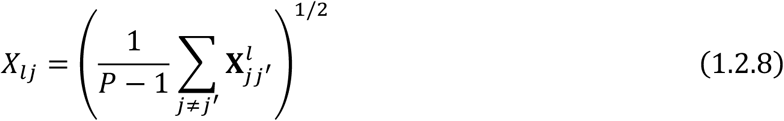

as the summary measure.

### STLs: Imputation for Cross-tissue Expression

We can use any predictive model as the learners in equation (1.2.1) to derive the effect size 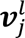. In this work, we used the combination of four different learners to demonstrate our TWAS-CTL method. We used penalized approaches such as LASSO (Least Absolute Shrinkage and Selection Operator, (Tibshirani, 1996)), Elastic Net (Zou & Hastie, 2005), and Ridge regression (Hoerl & Kennard, 1970). As recommended by (Gamazon et al., 2015), we also incorporated a machine learning methods (Hastie et al., 2009) -the random forest method (Breiman, 2001) into our framework. We employed a five-fold cross-validation to tune and evaluate the model.

### Gene-level Association Studies

The gene-trait association were detected through regression models based on traits and imputed gene expression. For example, for a continuous trait, the following regression model for a given gene 𝑔 was used:

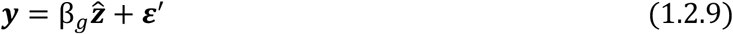

where, 𝒚 is the vector for trait, 𝛽_1_ is the total effect, 𝒛: is vector for the cross-tissue combined imputed gene expression, and 𝜺’ is the vector for errors. Finally, the gene-trait association can be detected by testing *H*_0_: β*_g_* = 0 vs *H*_1_: β*_g_* ≠ 0.

### Simulation Design

To demonstrate the performance of our proposed method, we used the genotype and phenotype data from Genetics of Kidneys in Diabetes (dbGaP accession id phs000018.v2.p1) in association studies and genotype data in the simulation studies. We applied to get the protected data from dbGaP from https://www.ncbi.nlm.nih.gov/projects/gap/cgi-bin/study.cgi?study_id=phs000018.v2.p1. The coordinates of genotype file depend on the human genome releases https://genome.ucsc.edu/FAQ/FAQreleases.html. Before we tested our method, we noted that the coordinates of genotype files depend on the specific version of the human genome used. Human genome releases represent different versions of the reference genome, and the genomic coordinates of a given SNP can vary between them. When a study was originally conducted, SNP coordinates were typically aligned to the latest genome release available at that time. Therefore, if the study was performed several years ago, the coordinates may not align with GRCh38, the current human genome build. Because many databases and computational tools rely on a standardized genome build, it is often necessary to convert SNP coordinates accordingly. We used the UCSC LiftOver tool (https://genome.ucsc.edu/cgi_bin/hgLiftOver) to convert SNP coordinates from GRCh36 to GRCh38. We applied the similar quality control that we applied to the GTEx genotype data.

The BiomaRt package from Bioconductor (ENSEMBL) provides a fast, user-friendly, and efficient interface for accessing BioMart directly within the R environment (Durinck et al., 2009). The GRCH37 annotation was used to allocate SNPs to the gene identifiers; this same annotation was also used to link gene IDs to gene names. We used Ensembl BioMart to assign SNPs to the corresponding gene. While some previous TWAS studies used 1-MB flank region to assign SNPs to a gene (Gamazon et al., 2015), recent studies using GTEx data suggest that the majority of cis-eQTLs are located within approximately 50–150 kb of the transcription start sites. For instance, GTEx v8 reports that >70% of the significant eQTLs fall inside the 100-kb flank region (Consortium et al., 2020). Hence, we used 100-kb flank region to assign SNPs to the genes.

We performed large-scale simulation to check the overall performance of TWAS-CTL, compared to UTMOST (Hu, Li, Lu, Weng, Wang, Zekavat, Yu, Li, Gu, Muchnik, et al., 2019). The simulation procedure which was followed in UTMOST has also been implied in our work. In the integration of diverse tissues, it is crucial to incorporate heterogeneity to recognize the variations among them, while also considering homogeneity where it might be applicable, especially among the tissues from the same organ. Therefore, in our simulation setting, for a given gene, we selected three tissues for our simulations: Nucleus Accumbens (𝑛 = 198), Brain Cortex (𝑛 = 204), and Esophagus Mucosa (𝑛 =492). These tissues exemplify unique biological systems, resulting in intrinsic heterogeneity owing to their varied cellular compositions and functions. Even though, same organ region may show significant variability due to their specialized functions, nonetheless, tissues from the same organ system (such as the Nucleus Accumbens and Brain Cortex) may also exhibit a degree of homogeneity in comparison to tissues from disparate systems, such as Esophagus Mucosa. For the i-th tissue, we generate the j-th random sample (trait): 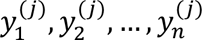 in three settings:

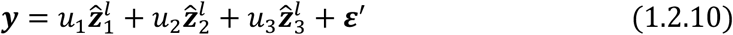

where, 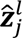 can be found using equation (1.2.2) - choosing imputed expression is critical to the simulation result. In contrast to the UTMOST study, which employed their method to obtain *̂v_j_* we used different predictive learning methods like Lasso, Ridge, Random Forest, PredixCan (Gamazon et al., 2015), and UTMOST to derive 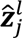 This approach opens windows of opportunity to see how the CTLs and UTMOST perform in diverse settings. On the other hand, 𝑢*_j_* ’s are the influences of the gene expression on the trait within the j-th tissue. To simulate data under the null hypothesis (type I error), we simply put 𝑢_1_ = 𝑢_2_ = 𝑢_3_ = 0.

To obtain power through simulation, we use a varied configuration that encompasses the characteristics of distinct tissues from both same and different organs. In the first configuration, we characterized it as 𝑢_1_ dominant, representing a less diversified environment (homogeneity) where the effect is concentrated in a single tissue (Nucleus Accumbens), hence decreasing the variability introduced by other tissues. This context presupposes homogeneity, as a single tissue is affecting the phenotype. In the second configuration, we implemented moderate heterogeneity (setting 𝑢_1_ = 𝑢_2_ = 𝑢_3_ = 0), resulting in an equitable distribution of effects between two distinct tissues (Nucleus Accumbens and Esophagus Mucosa), hence augmenting variability relative to the initial setup. In the third configuration, we assign 𝑢_1_ = 𝑢_2_ = 𝑢_3_ = 1/3, resulting in a uniform effect across all three tissues, illustrating the true characteristics of various tissues from the same and distinct organs. This configuration signifies the highest degree of heterogeneity, as all chosen tissues contribute equally to the phenotype.

We used the genotype information from the GoKinD data as 𝐆_’_ (as in the equation 1.2.2) in our simulation setup. We then performed extensive simulations after we implemented the TWAS-CTL method to estimate cross-tissue gene-trait association- with varying sample size of 500 and 750, replicating the whole process 1000, 5000, and 10000 times for a given gene. Statistical significance 0.05 was used as a threshold to calculate the power of the test.

## RESULTS

### Simulation Studies: Empirical Type I Error Rate

We compared the simulation results of our method with UTMOST (implemented as described in (Hu, Li, Lu, Weng, Wang, Zekavat, Yu, Li, Gu, Muchnik, et al., 2019; Zhou et al., 2020)). In addition, we implemented PrediXcan (Gamazon et al., 2015) to impute 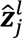 to compare our method with UTMOST, where we implemented Generalized Berk-Jones (GBJ) test (Sun & Lin, 2017) to combine the imputed gene expression from different tissues as UTMOST did. To assess type I error control, we compared TWAS-CTL (averaged across six TWAS-CTLs: LASSO+Ridge, LASSO+Elastic Net, LASSO+Random Forest, Elastic Net+Ridge, Elastic Net+Random Forest, and Ridge+Random Forest) with UTMOST across three sample sizes (𝑛 = 500, 750, and 1000) and three replication levels (𝑟 = 1000, 5000, and 10,000). As shown in **Table 1**, both methods maintained the type I error rate close to the nominal threshold of α = 0.5; however, TWAS-CTL consistently achieved slightly more conservative control under high-replication settings. At 𝑛 = 500, UTMOST exhibited an inflated error rate of 0.0620 when r = 1,000, while TWAS-CTL remained well-controlled at 0.0440. As the number of replications increased, both methods converged, but TWAS-CTL consistently exhibited a lower type I error (0.0474) compared to UTMOST (0.0519) at 𝑟 = 10,000. A similar trend was observed at the sample size of 𝑛 = 750, where TWAS-CTL achieved lower or comparable type I error rates relatively to UTMOST. At the largest sample size (𝑛 = 1000), TWAS-CTL showed modest inflation at 𝑟 = 1,000; however, this was corrected as the number of replications increased,, ultimately producing the lowest type I error under the most reliable conditions. Collectively, these findings suggest that TWAS-CTL offers consistently strong control of false positives, particularly under adequate replication, underscoring its robustness in simulation-based hypothesis testing.

**Table 1.** Type I error rates for TWAS-CTL and UTMOST across different sample sizes (𝑛) and replications (𝑟).

| $N$ | $M$ | TWAS-CTL | UTMOST |
| --- | --- | --- | --- |
| **500** | **1000** | 0.0440 | 0.0620 |
|  | **5000** | 0.0516 | 0.0518 |
|  | **10000** | 0.0474 | 0.0519 |
| **750** | **1000** | 0.0470 | 0.0480 |
|  | **5000** | 0.0490 | 0.0482 |
|  | **10000** | 0.0476 | 0.0491 |
| **1000** | **1000** | 0.0570 | 0.0460 |
|  | **5000** | 0.0492 | 0.0522 |
|  | **10000** | 0.0476 | 0.0485 |

### Simulation Studies: Statistical Power

**Table 2** summarizes the statistical power of six TWAS-CTL methods and UTMOST under three settings of tissue-specific genetic influence settings: (1, 0, 0), (1, 1, 0), and (1/3, 1/3, 1/3), corresponding to U settings 1, 2, and 3, respectively. Simulations were conducted with a sample size of *n=*750 and 10,000 replications. Predicted expression values, denoted as 𝒛:^$^ , were imputed using the random forest approach. The methods compared include various CTL combinations – Enet + RF (Elastic Net + Random Forest), Enet + Ridge (Elastic Net + Ridge Regression), Lasso + RF (Lasso + Random Forest), Lasso + Ridge (Lasso + Ridge Regression), Ridge + RF (Ridge Regression + Random Forest) - and UT (UTMOST). **Figure 3** illustrates that Enet + Ridge and Lasso + Ridge CTLs demonstrated consistently high power in both homogeneous and somewhat heterogeneous conditions, indicating that these approaches are robust across varying levels of tissue-specific effects. In contrast, Lasso + RF showed perfect power in moderate and highly heterogeneous settings, establishing it as the most potent method in these scenarios. UTMOST also demonstrated significant power in environments characterized by moderate and high heterogeneity. These findings indicated that Lasso + RF CTL was very effective in heterogeneous setting, whereas Enet + Ridge and Lasso + Ridge CTLs were effective in homogeneous and somewhat diverse contexts.

**Figure 3.**
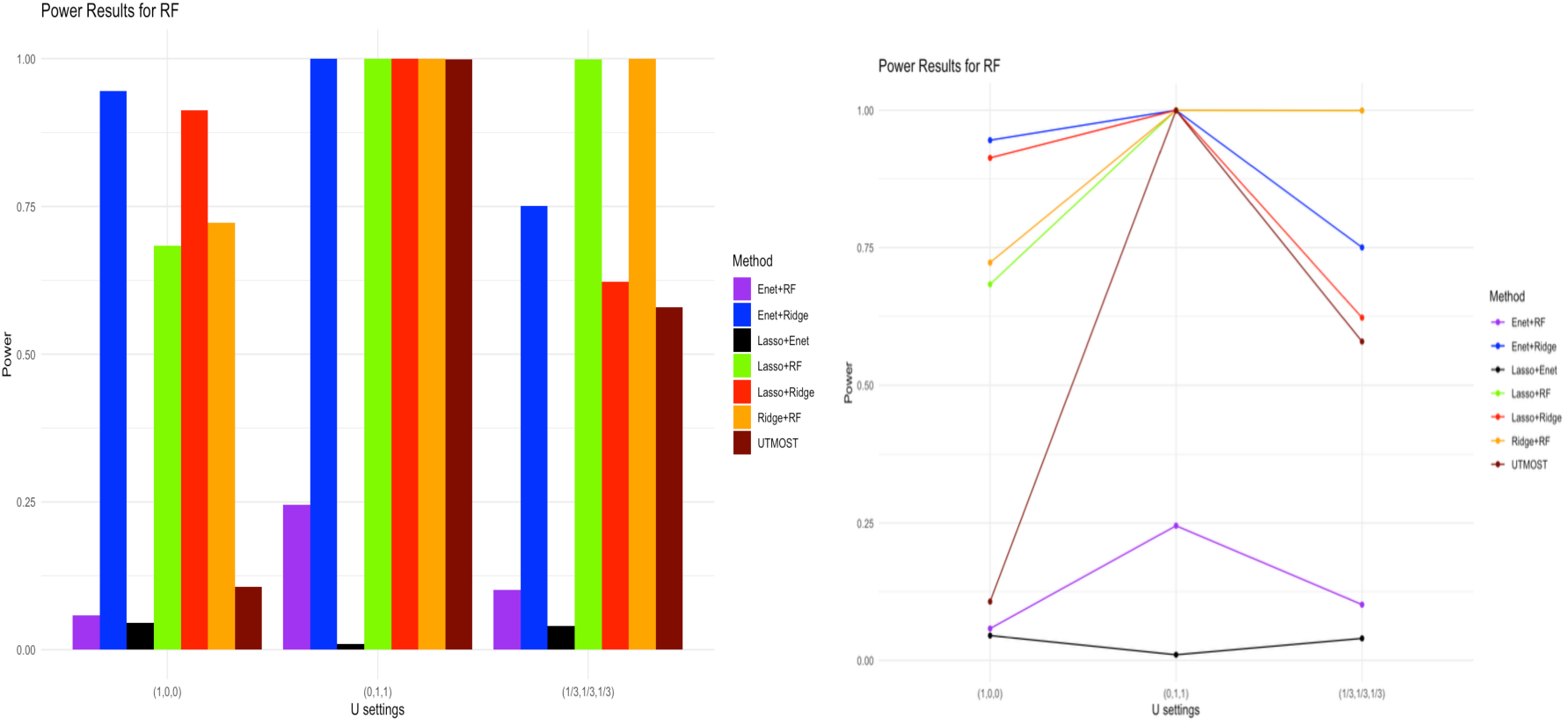
Visualization of power comparison between different TWAS-CTLs and UTMOST when the random forest method was used as the imputed expressions (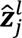).

**Table 2.** Power of TWAS-CTL combinations and UTMOST across different levels of cross-tissue heterogeneity with the sample size of 𝑛 = 750 and different number of replicates (𝑟) when the random forest method was used to impute 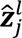. The highest power for 10,000 replications in each setting is bold.

| $U$ | $r$ | E + Ri | L + Ri | L + E | L + RF | E + RF | Ri + RF | UT |
| --- | --- | --- | --- | --- | --- | --- | --- | --- |
| **1** | 1000 | 0.9490 | 0.9050 | 0.0340 | 0.6750 | 0.0560 | 0.7240 | 0.1040 |
|  | 10000 | **0.9454** | 0.9132 | 0.0451 | 0.6833 | 0.0578 | 0.7227 | 0.1069 |
| **2** | 1000 | 1.0000 | 1.0000 | 0.0100 | 1.0000 | 0.2470 | 1.0000 | 1.0000 |
|  | 10000 | **1.0000** | **1.0000** | 0.0101 | **1.0000** | 0.2450 | **1.0000** | 0.9994 |
| **3** | 1000 | 0.7450 | 0.6150 | 0.0490 | 1.0000 | 0.1280 | 1.0000 | 0.5640 |
|  | 10000 | 0.7503 | 0.6226 | 0.0399 | 0.9994 | 0.1014 | **0.9996** | 0.5791 |

Enet + Ridge, Lasso + Ridge, and Enet + RF CTLs consistently demonstrated strong robustness across varying degrees of tissue influence. When LASSO was used to impute 𝒛:^$^, Ridge + RF CTL also performed well, particularly under moderately heterogeneous settings. In contrast, UTMOST generally exhibited diminished power across all configurations relative to the other methods (**Table 3**). As shown in **Figure 4**, Enet + Ridge, Lasso + Ridge, and Enet + RF CTLs maintained very high power in homogeneous and moderately heterogeneous settings, and high power even under highly heterogeneous conditions. These methods were consistently reliable across different degrees of tissue influence. Ridge + RF CTL achieved high power in moderately heterogeneous settings and moderate power in homogeneous and highly heterogeneous settings, whereas Lasso + RF CTL showed moderate power in homogeneous and moderately heterogeneous settings. UTMOST displayed generally lower power across all settings compared with the other methods.

**Figure 4.**
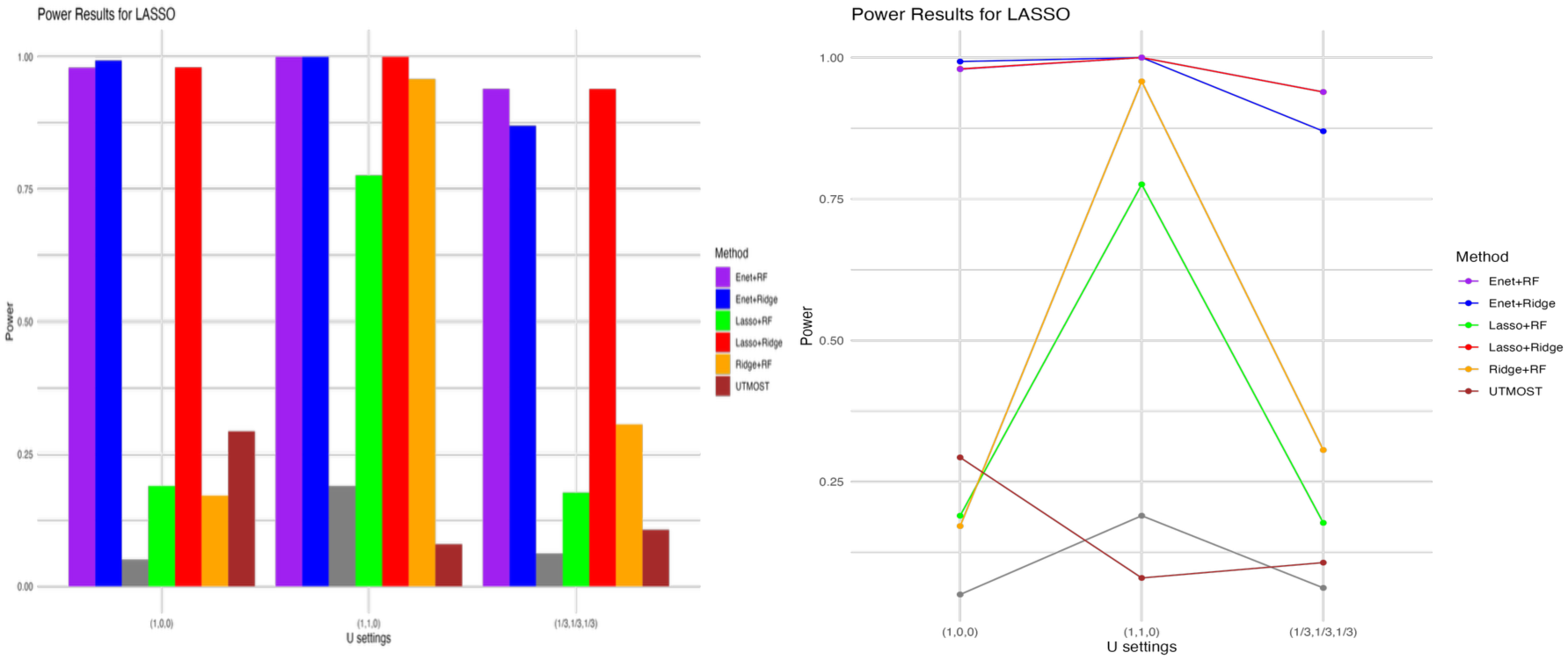
Visualization of power comparison between different TWAS-CTLs and UTMOST when LASSO was used as the imputed expressions (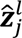).

**Table 3.** Power of TWAS-CTL combinations and UTMOST across varying levels of cross-tissue heterogeneity with the sample size of 𝑛 = 750 and different number of replicates (𝑟) when LASSO was used to impute 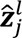. The highest power for 10,000 replications in each setting is bold.

| $U$ | $r$ | E + Ri | L + Ri | L + E | L + RF | E + RF | Ri + RF | UT |
| --- | --- | --- | --- | --- | --- | --- | --- | --- |
| **1** | 1000 | 0.9930 | 0.9770 | 0.0470 | 0.1890 | 0.9750 | 0.1530 | 0.2400 |
|  | 10000 | **0.9930** | 0.9800 | 0.0506 | 0.1899 | 0.9794 | 0.1716 | 0.2930 |
| **2** | 1000 | 1.0000 | 1.0000 | 0.1940 | 0.7860 | 1.0000 | 0.9680 | 0.0680 |
|  | 10000 | **1.0000** | **1.0000** | 0.1899 | 0.7758 | **1.0000** | 0.9578 | 0.0221 |
| **3** | 1000 | 0.8700 | 0.9380 | 0.0600 | 0.1860 | 0.9350 | 0.2840 | 0.1000 |
|  | 10000 | 0.8698 | 0.9391 | 0.0623 | 0.1773 | **0.9392** | 0.3061 | 0.1070 |

When the ridge regression was used as the imputation method (**Table 4**), Enet + Ridge and Lasso + Ridge CTLs exhibited strong power across varying degrees of tissue heterogeneity settings, Ridge + RF also performed well, particularly in moderately heterogeneous settings, whereas UTMOST showed comparatively low power in all settings, achieving only moderate power under heterogeneity. As shown in **Figure 5**, Enet + Ridge and Lasso + Ridge achieved very high power in homogeneous settings and moderate to high power under heterogeneous conditions, demonstrating consistent reliability across different levels of tissue influence. Ridge + RF performed best in moderately heterogeneous settings and maintained high power in homogeneous settings while Lasso + RF displayed moderate power in both homogeneous and moderately heterogeneous settings. In contrast, UTMOST consistently exhibited low power than the other methods, except under moderate heterogeneity, where its performance improved modestly. Overall, these results suggest that Enet + Ridge and Lasso + Ridge are consistently powerful and reliable across a range of tissue influence settings, while Ridge + RF also show strong performance, particularly in moderately heterogeneous conditions.

**Figure 5.**
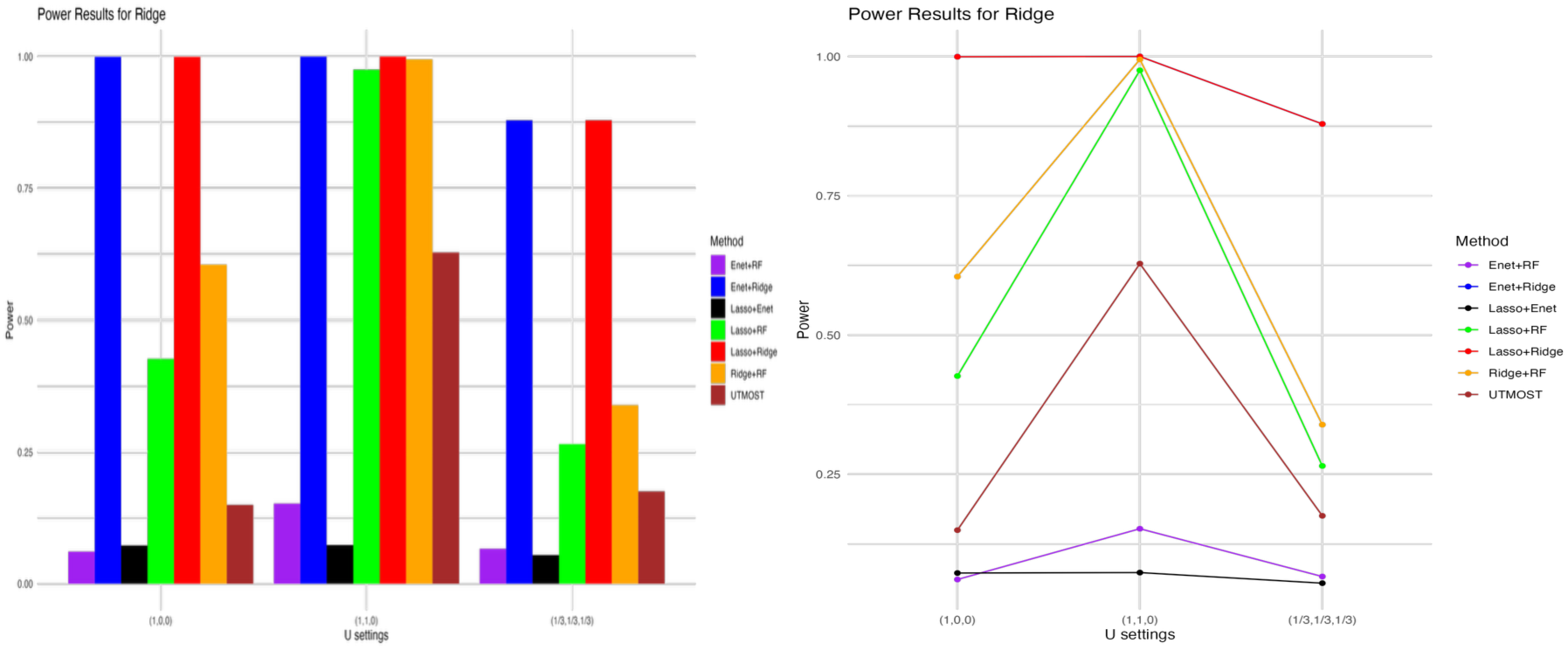
Visualization of power comparison between different TWAS-CTLs and UTMOST when the ridge regression was used as the imputed expressions (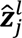).

**Table 4.** Power of TWAS-CTL combinations and UTMOST across varying levels of cross-tissue heterogeneity with the sample size of 𝑛 = 750 and different number of replicates (𝑟) when the ridge regression was used to impute 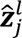. The highest power for 10,000 replications in each setting is bold.

| $U$ | $r$ | **E + Ri** | **L + Ri** | **L + E** | **L + RF** | **E + RF** | **Ri + RF** | **UT** |
| --- | --- | --- | --- | --- | --- | --- | --- | --- |
| **1** | 1000 | 1.0000 | 0.9990 | 0.0670 | 0.4150 | 0.0480 | 0.5920 | 0.1360 |
|  | 10000 | **0.9996** | 0.9995 | 0.0728 | 0.4264 | 0.0612 | 0.6049 | 0.1498 |
| **2** | 1000 | 1.0000 | 1.0000 | 0.0900 | 0.9740 | 0.1510 | 0.9950 | 0.7000 |
|  | 10000 | **1.0000** | **1.0000** | 0.0736 | 0.9751 | 0.1523 | 0.9947 | 0.3983 |
| **3** | 1000 | 0.8810 | 0.8805 | 0.0610 | 0.2630 | 0.0810 | 0.3420 | 0.1750 |
|  | 10000 | **0.8789** | 0.8786 | 0.0546 | 0.2650 | 0.0665 | 0.3391 | 0.1787 |

As UTMOST was used to obtain the imputed expression (**Table 5**), it consistently showed high or even perfect power across all settings, making it the most powerful method in different settings. In comparison, Lasso + RF and Ridge + RF CTLs showed moderate power under moderately and highly heterogeneous conditions.

**Table 5:** Power comparison of TWAS-CTL combinations and UTMOST across varying levels of cross-tissue heterogeneity (sample size 750, m replications) when UTMOST was used to impute 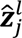. The highest power for 10,000 replications in each U setting is bold.

| $U$ | $r$ | E + Ri | L + Ri | L + E | L + RF | E + RF | Ri + RF | UT |
| --- | --- | --- | --- | --- | --- | --- | --- | --- |
| **1** | 1000 | 0.0580 | 0.1100 | 0.0370 | 0.1020 | 0.0330 | 0.0900 | 0.9850 |
|  | 10000 | 0.0561 | 0.1076 | 0.0476 | 0.0967 | 0.0471 | 0.0904 | **0.9849** |
| **2** | 1000 | 0.0370 | 0.1720 | 0.0450 | 0.2820 | 0.0510 | 0.2620 | 1.0000 |
|  | 10000 | 0.0376 | 0.1711 | 0.0344 | 0.2832 | 0.0361 | 0.2559 | **1.0000** |
| **3** | 1000 | 0.0490 | 0.0900 | 0.0620 | 0.1050 | 0.0580 | 0.1000 | 0.9988 |
|  | 10000 | 0.0491 | 0.0879 | 0.0487 | 0.1124 | 0.0503 | 0.1036 | **0.9985** |

In the comparative simulation utilizing various imputation methods for expression (𝒛:^$^), it was observed that when PrediXcan was employed (**Table 6**), Enet + Ridge and Lasso + Ridge CTLs were consistently reliable across scenarios with varying tissue influences, whereas UTMOST offered a consistently moderate power instead. As shown in **Figure 6**, Enet + Ridge and Lasso + Ridge CTLs demonstrated high power under both homogeneous and highly heterogeneous settings. Under moderate heterogeneity, Ridge + RF CTL and UTMOST exhibited comparable power. Overall, UTMOST maintained moderate power across different settings, outperforming Lasso + Enet, Lasso + RF, and Enet + RF CTLs.

**Figure 6.**
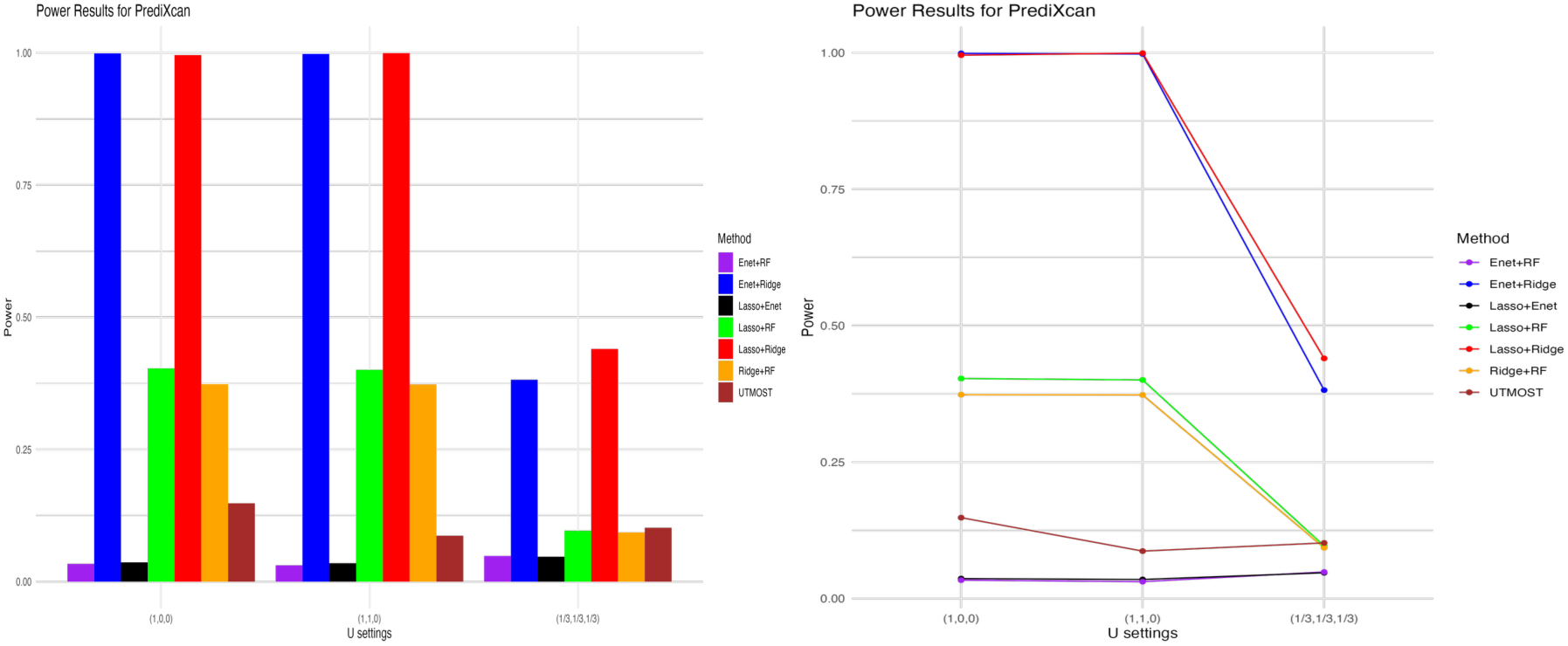
Visualization of power comparison between different TWAS-CTLs and UTMOST when PrediXcan was used as the imputed expressions (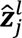).

**Table 6.** Power of TWAS-CTL combinations and UTMOST across varying levels of cross-tissue heterogeneity with the sample size of 𝑛 = 750 and different number of replicates (𝑟) when PrediXcan was used to impute 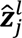. The highest power for 10,000 replications in each setting is bold.

| $U$ | $r$ | E + Ri | L + Ri | L + E | L + RF | E + RF | Ri + RF | UT |
| --- | --- | --- | --- | --- | --- | --- | --- | --- |
| **1** | 1000 | 0.9990 | 0.9960 | 0.0310 | 0.4040 | 0.0190 | 0.3760 | 0.1390 |
|  | 10000 | **0.9988** | 0.9956 | 0.0365 | 0.4031 | 0.0337 | 0.3735 | 0.1482 |
| **2** | 1000 | 1.0000 | 1.0000 | 0.0470 | 0.3890 | 0.0430 | 0.3700 | 0.0960 |
|  | 10000 | 0.9978 | **0.9992** | 0.0349 | 0.4005 | 0.0309 | 0.3730 | 0.0869 |
| **3** | 1000 | 0.3900 | 0.4560 | 0.0610 | 0.0850 | 0.0610 | 0.0790 | 0.1090 |
|  | 10000 | 0.3818 | **0.4400** | 0.0472 | 0.0963 | 0.0486 | 0.0931 | 0.1019 |

As shown in **Figure 7**, Enet + Ridge and Lasso + Ridge CTLs consistently demonstrated higher power across all settings, indicating that these combinations are robust and perform reliably well regardless of data heterogeneity. This consistency makes them strong choices for CTL analysis in diverse tissue scenarios. Although power was generally lower under homogeneity conditions, Ridge + RF CTL and UTMOST showed promising performance in in moderately and highly heterogeneous settings, reflecting their ability to adapt to data variability. Overall, the analysis highlights the complementary strengths of different methods. Enet + Ridge and Lasso + Ridge CTLs exhibited robustness and versatility across all scenarios**, while** Ridge + RF and UTMOST performed particularly well in moderate and high heterogeneous settings, leveraging their compacity to handle both linear and non-linear interactions. UTMOST ’s strong performance under moderate to high heterogeneity further validates its design for multi-tissue modeling.

**Figure 7.**
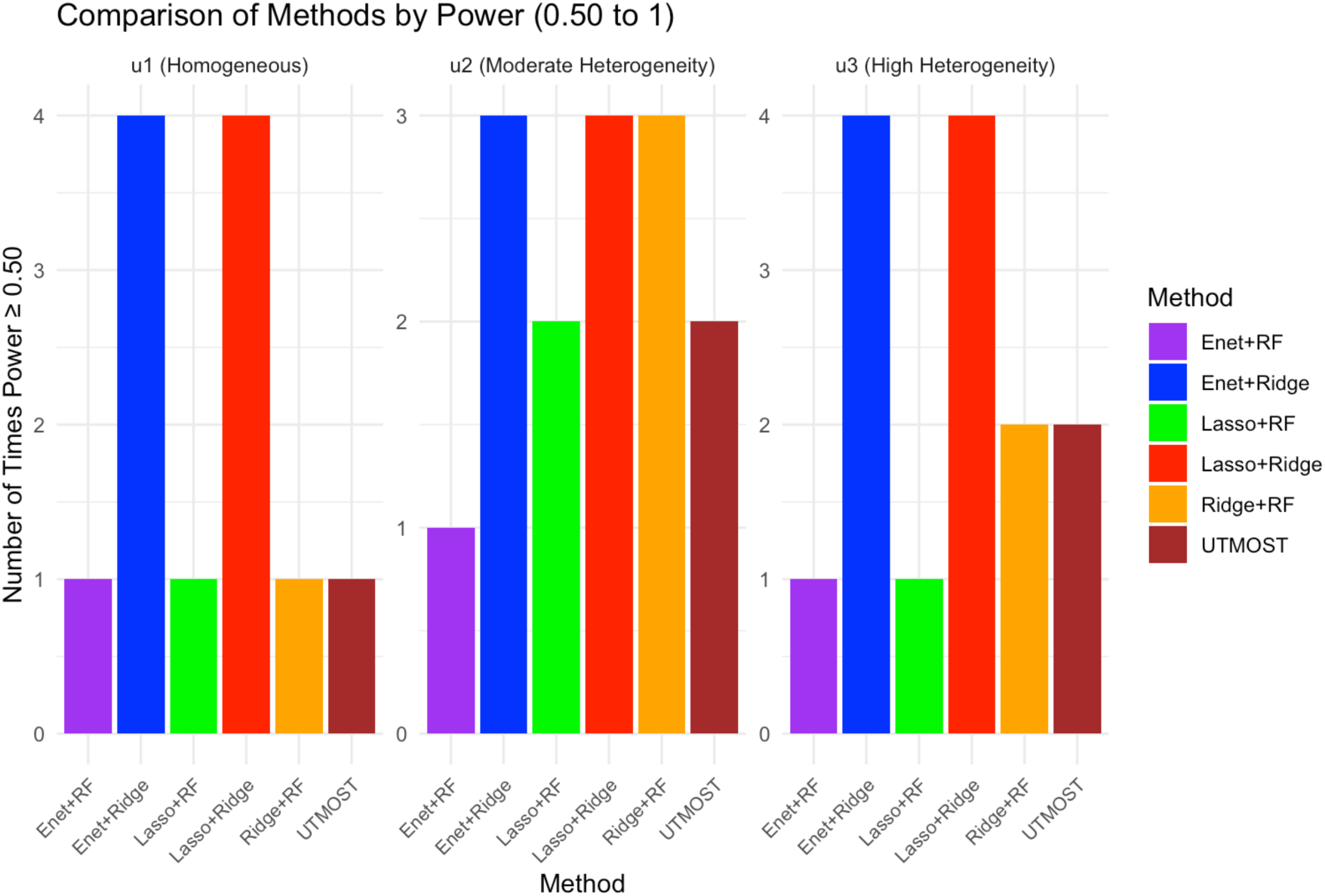
Comparison of different TWAS-CTLs and UTMOST by power across three levels of heterogeneity - the methods compared are: the Y-axis consists of different TWAS-CTLs- Enet+RF (Elastic Net + Random Forest), Enet+Ridge (Elastic Net + Ridge Regression), Lasso+RF (Lasso + Random Forest), Lasso+Ridge (Lasso + Ridge Regression), Ridge+RF (Ridge Regression + Random Forest), and UTMOST. The X-axis indicates how many times the power was 0.5 or more across different methods of imputation to get the imputed expression (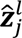).

### Application to Real Data

As an illustration of our proposed method, we applied it along with UTMOST, and PrediXcan to the Genetics of Kidneys in Diabetes (GoKinD) cohort, an association study for type I diabetes. After quality control, we retained 8859 genes and 62701 common SNPs in the GoKinD data across 1108 individuals. Accordingly, the genome-wide significance threshold after applying the Bonferroni correction is 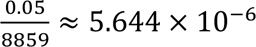. Nevertheless, Bonferroni correction can increase false negatives rate in multi-tissue TWAS leading to overly conservative thresholds and reduced sensitivity to true effects (Alvaro N Barbeira et al., 2019). In previous studies, a 𝑝-value threshold for significance of 1 × 10^−4^ was widely used for gene based association testing (Fischer et al., 2018; Gao et al., 2021). It is important to note that GoKinD data has relatively small sample size compared to other typical GWAS datasets, which often include millions of individuals with thousands of genetic variants linked to the genes. Henceforth, we used the threshold of 3 × 10^−4^ to report the significant genes - which may serve as a balance between discovery and false-positive control when examining genome-wide expression-trait associations in a dataset of comparatively smaller sample size. As described before, type 1 diabetes was the phenotype of interest in this analysis.

**Table 7** highlights a set of genes identified as statistically significant using the TWAS-CTL framework. Each gene listed in the table has been previously implicated in various complex traits or diseases, as documented in the GWAS Catalog (https://www.ebi.ac.uk/gwas/home). The “Description” column in **Table 7** summarizes these associations, providing valuable biological context for their relevance to type I diabetes. Notably, FCHSD2 (11:72836745–73142318) surpassed the Bonferroni-corrected genome-wide significance threshold, showing the most significant p-value in the analysis (2.81 × 10^−6^). This gene has been linked to several autoimmune and metabolic traits related to type-1 diabetes, including systemic lupus erythematosus (Su et al., 2023), Crohn’s disease (Chranioti et al., 2022), Psoriasis (Abramczyk et al., 2020), HbA1c levels (Patiño-Fernández et al., 2009). Such pleiotropy underscores its potential regulatory role in immune–metabolic crosstalk. Similarly, BFSP2 (3:133400056-133475222) and KLHL8 (4:87160103-87240314), both genes identified with the p-value of 8.72 × 10^−6^ *and* 2.81 × 10^−5^ respectively, were also associated with traits relevant to type 1 diabetes. These includes diffuse plaque measurement (Orchard et al., 2006), serum alanine aminotransferase measurement (West et al., 2006), platelet count (Schneider, 2009), hemoglobin A1c (HbA1c) testing (Eyth & Naik, 2023), and total cholesterol measurement (Candler et al., 2017). Collectively, these findings reinforce the biological plausibility of the TWAS-CTL results and highlight genes involved in metabolic and inflammatory pathways linked to type 1 diabetes.

**Table 7.** Significant genes identified by TWAS-CTL at a p-value threshold 3 × 10^−4^. Gene-related trait information listed in the “Description” column was obtained from the GWAS Catalog, indicating phenotypes previously associated with these genes. Citations within the description denote prior studies linking these traits to type 1 diabetes. Note that genes with Bonferroni corrected threshold setting are bolded and new genes are being bold and italic.

| Gene<br>*p*-value | Description |
| --- | --- |
| **FCHSD2**<br>$2.81 \times 10^{-6}$ | Systemic lupus erythematosus (Su et al., 2023), Crohn's disease (Chranioti et al., 2022), Psoriasis (Abramczyk et al., 2020), HbA1c measurement (Patiño-Fernández et al., 2009), etc. |
| BFSP2<br>$8.72 \times 10^{-6}$ | Diffuse plaque measurement (Orchard et al., 2006), Alzheimer's disease (Lacy et al., 2018), Gut microbiome (Zheng et al., 2018), etc. |
| KLHL8<br>$1.94 \times 10^{-5}$ | Serum Alanine Aminotransferase measurement (West et al., 2006), Platelet count (Schneider, 2009), Hemoglobin A1c (HbA1c) testing (Eyth & Naik, 2023), Total Cholesterol measurement (Candler et al., 2017), etc. |
| RFC5<br>$5.25 \times 10^{-5}$ | Neuroimaging measurements (Filip et al., 2020), type 2 diabetes (Bielka et al., 2024), IGFBP-1 measurement (Sharma et al., 2016), Systemic Lupus Erythematosus measurement (Su et al., 2023), etc. |
| VKORC1L1<br>$5.53 \times 10^{-5}$ | Glomerular filtration rate (Group, 2011), Neutrophil measurement and count (Huang et al., 2016), DPP4 protein level (Wang et al., 2018), etc. |
| MYO3B<br>$6.61 \times 10^{-5}$ | Retinal vasculature measurement (Sousa et al., 2020), Lipid measurement (Nelson et al., 2020), Serum protein measurement (Zhi et al., 2011), etc. |
| ZC3H18<br>$7.12 \times 10^{-5}$ | Erythrocyte volume (Wang et al., 2021), Platelet count (Schneider, 2009), Aspartate aminotransferase measurement (Asada et al., 2018), C-peptide measurement (Maddaloni et al., 2022), etc. |
| EPHA4<br>$1.14 \times 10^{-4}$ | Protein measurement (Zhi et al., 2011), Cerebral cortex area attribute (Lyoo et al., 2013), PHF-tau measurement (Hobday et al., 2021), etc. |
| UCHL1<br>$1.70 \times 10^{-4}$ | Diabetic mastopathy (Boumarah et al., 2022), HDL cholesterol levels (Kothari et al., 2024), Bone mineral density mean (Loxton et al., 2021), etc. |
| ME3<br>$1.99 \times 10^{-4}$ | Erythrocyte count (Herance et al., 2023), Open-angle glaucoma (Zhou et al., 2014), Lymphocyte count (Miao et al., 2008), Chronic obstructive pulmonary disease (Park et al., 2022), etc. |
| USP8<br>$2.17 \times 10^{-4}$ | Monocyte count (Bradshaw et al., 2009), C-reactive protein levels (Kouchaksaraei et al., 2022), Interleukin-6 (IL-6) levels (Gomes, 2017), etc. |
| TGS1<br>$2.22 \times 10^{-4}$ | Neutrophil count (Huang et al., 2016), leukocyte quantity (Menart-Houtermans et al., 2014), monocyte count (Bradshaw et al., 2009), platelet component distribution width (Schneider, 2009), etc. |
| SERINC1<br>$2.47 \times 10^{-4}$ | Plateletcrit (Schneider, 2009), heat shock protein beta-1 levels, monocyte count (Moin et al., 2021), lymphocyte count (Miao et al., 2008), etc. |
| ***RP11-550H2***<br>$2.90 \times 10^{-4}$ | New Gene |
| RAPGEF1<br>$2.91 \times 10^{-4}$ | Gut microbiome measurement (Zheng et al., 2018), Thyroid stimulating hormone (Umpierrez et al., 2003), Hypertrophic cardiomyopathy (Lee et al., 2022), etc. |

Most importantly, using the proposed significance threshold, TWAS-CTL identified a previously unreported gene, Long Intergenic non-protein Coding RNA (RP11-550H2, located at chr1:76,507,376–76,531,913), with a derived p-value of 2.90 × 10^−4^. This discovery suggests that RP11-550H2 may represent a previously unrecognized contributor to disease biology, warranting further investigation through functional validation.

In total, TWAS-CTL identified 15 significant genes (**Table 7**), whereas UTMOST identified 2 genes- MAST2 (1:45786987-46036122, p-value: 2 × 10^-5^) and LINC00575 (4:82610974-82621610, p-value: 2.13 × 10-5) in our suggested significance threshold. In comparison, PrediXcan (Gamazon et al., 2015), which combines the cross-tissue imputed gene expressions with the Generalized Berk-Jones (GBJ) test (Sun & Lin, 2017) as implemented in UTMOST, identified a single gene–KLHL8 (p-value: 9.37 × 10^−5^). Notably, KLHL8 was also identified by TWAS-CTL, but with a more stringent and higher significant 𝑝-value of 2.81 × 10^−6^. When a more relaxed significance threshold of 1 × 10^−3^ was used, **TWAS-CTL** identified **35 genes**, whereas **UTMOST** and **PrediXcan** detected only **4** and **8 genes**, respectively. These results demonstrate that TWAS-CTL achieves a stronger balance between sensitivity and specificity, capturing more biologically relevant associations while maintaining rigorous statistical control.

## DISCUSSION

Transcriptome-wide association studies (TWAS) leverage the genetic component of gene expression to elucidate gene-trait mechanisms. However, most multi-tissue TWAS methods combine association results only after conducting single-tissue tests, typically by aggregating their 𝑝-values using procedures such as the Generalized Berk-Jones statistic. In contrast, TWAS-CTL (Cross-Tissue Learner) introduces an early-fusion strategy: immediately after imputation, predicted expression profiles from multiple tissues are integrated through an adaptive weighting function that amplifies informative tissues and attenuates noisier ones. By incorporating cross-tissue covariance at the modelling stage, TWAS-CTL preserves inter- and intra-tissue regulatory information, reduces the multiple-testing burden, and is designed to enhance statistical power while maintaining proper control of type-I errors.

We compared TWAS-CTL with two commonly used methods - using extensive simulations that span homogeneous to highly heterogeneous tissue architectures (UTMOST), as well real data from the Genetics of Kidneys in Diabetes (GoKinD) cohort. In the simulation studies assessing type I error rate, across four sample sizes (𝑛 = 1000, 2000, 5000, and 10000) and three replication levels (𝑟 = 1000, 5000, 10000), TWAS-CTL consistently maintained the nominal α = 0.05 rate. In contrast, UTMOST exhibited inflated type I error at the smallest sample 𝑛 = 1000 and lowest replication 𝑟 = 1,000. Although both methods converged with increasing Monte-Carlo iterations (Metropolis & Ulam, 1949), TWAS-CTL remained slightly more conservative, especially when 𝑟 = 10,000. These findings imply that the TWAS-CTL weighting mechanism guards against spurious associations even when training data are sparse or when the simulation space is undersampled, a common limitation in high-dimensional genomic studies.

Power analyses were structured around three tissue-weight configurations representing mimic homogeneous, moderately heterogeneous, and highly heterogeneous genetic architectures. When expressions were imputed with the random forest method, CTL configurations pairing a linear learner with Ridge regression (Enet + Ridge; Lasso + Ridge) were consistently powerful, across all heterogeneity levels. By contrast, CTLs employing random forest method in the second stage (e.g., Lasso + RF) excelled only when tissue effects diverged strongly, highlighting their suitablity for non-linear or interaction-dominated contexts. These patterns persisted when the imputation engine was switched to Elastic Net, Lasso, or Ridge alone: the Enet + Ridge and Lasso + Ridge pairings again achieved superior power, underscoring the advantages of a hybrid linear-penalized strategy in balancing bias and variance. UTMOST, while competitive in moderate heterogeneity, underperformed in homogeneous and highly heterogeneous settings, suggesting that its single-model framework struggles when tissue contributions are either uniform or extremely discordant.

Applying TWAS-CTL to the GoKinD cohort – comprising 8,859 genes interrogated with approximately 63, 000 common SNPs in 1,108 individuals - yielded 15 genome-wide signals at our pragmatic significance threshold of 𝑝 − 𝑣𝑎𝑙𝑢𝑒 < 3 × 10^−4^. Notably, FCHSD2 exceeded even the conservative Bonferroni cut-off, and its prior links to systemic lupus erythematosus, Crohn’s disease, psoriasis, and glycated-haemoglobin levels provide a biologically coherent bridge to type 1 diabetes. Each additional gene identified by TWAS-CTL was likewise enriched for immune-metabolic traits overlapping the type 1 diabetes etiological spectrum, underscoring the method’s ability to priorities mechanistically plausible loci. Perhaps most compelling is the discovery of the long intergenic non-coding RNA **RP11-550H2**, a novel locus whose emergence illustrates the key advantage of early cross-tissue aggregation: distributed yet concordant expression signals that are too weak to reach significance in any single tissue become detectable when modeled jointly. Perhaps most compelling is the discovery of the long intergenic non-coding RNA RP11-550H2, a novel locus whose emergence illustrates the key advantage of early cross-tissue aggregation: distributed yet concordant expression signals that are too weak to achieve significance in any single tissue become detectable when modeled jointly. Methodological comparisons reinforce this conclusion. Under the same significance criterion, UTMOST identified only two genes and PrediXcan a single gene. Even when the threshold was relaxed to 1 × 10^<6^, TWAS-CTL retrieved 35 genes, whereas UTMOST and PrediXcan recovered just four and eight, respectively. Thus, TWAS-CTL is not simply more permissive; rather, it concentrates true biological signal more efficiently by integrating complementary tissue information at the modelling stage.

From a computational standpoint, TWAS-CTL is substantially efficient than UTMOST. All simulation experiments were conducted on a Dell Inspiron (13-th Gen Intel Core i5, 8 GB RAM, DDR5). Across all learner configurations, TWAS-CTL completed its runs in ≤ 50 % of the wall-clock time required by UTMOST under identical input dimensions. The difference was even more pronounced in real-data analysis. Using a MacBook pro (Apple M3 max, 64 GB memory, 16 core CPU, 40 core GPU), TWAS-CTL finished the full GoKinD scan in approximately 7–8 h for penalized-regression ensembles (LASSO, Elastic Net, Ridge) and ≈ 15–17 h when Random Forest was included. By contrast, the UTMOST pipeline–powered by sparse-group LASSO–required nearly seven days to finish the same dataset. Similar extremely prolonged runtimes for UTMOST have been reported previously (Morgante et al., 2023), underscoring its scalability limitations. Collectively, these results demonstrated that TWAS-CTL is not only statistically robust but also computationally practical for large-scale, multi-tissue TWAS applications.

Despite its strengths, multi-tissue TWAS presents several limitations that warrant further investigation. First, cross-study reproducibility remains imperfect: a gene–trait signal that is robust in one GWAS may attenuate or even disappear in another due to differences in allele frequencies, LD patterns or ancestry composition. To address this, our next iteration aims to incorporate cohort-specific GWAS information into the cross-tissue weighting step, allowing tissue weights to reflect both expression predictability and trait-specific SNP architecture. Additionally, training multiple learner pairs across 49 tissues is computationally intensive. Future work might explore dimensionality reduction in tissue space to improve scalability. Finally, the GoKinD dataset is modest by today’s standards. A critical next step is to deploy TWAS-CTL on biobank-scale resources such as the UK Biobank; which provides both the statistical power and ancestral diversity needed to validate novel loci and refine tissue-specific regulatory maps.

In summary, from the architecture of TWAS-CTL to its final data analysis and extensive simulation, four key themes emerged: (i) TWAS-CTL consistently maintained nominal type I error rates, (ii) it reached higher or at least equivalent statistical power across a wide range of tissue heterogeneity, (iii) it identified a broader set of biologically plausible and interpretable gene–trait associations even in a moderately sized case–control samples, and (iv) it supports plug-and-play predictive learners and user-defined utility functions. TWAS-CT’s early fusion strategy captures shared regulatory signal across tissues while preserving the tissue-specific effects critical to organ-focused pathophysiology. By balancing cross-tissue commonalities with biological heterogeneity, TWAS-CTL offers a robust and efficient framework for multi-tissue TWAS and is well positioned for application in large-scale studies.

